# Learning Nonlinear Brain Dynamics: van der Pol Meets LSTM

**DOI:** 10.1101/330548

**Authors:** Germán Abrevaya, Aleksandr Aravkin, Guillermo Cecchi, Irina Rish, Pablo Polosecki, Peng Zheng, Silvina Ponce Dawson

## Abstract

Many real-world data sets, especially in biology, are produced by highly multivariate and nonlinear complex dynamical systems. In this paper, we focus on brain imaging data, including both calcium imaging and functional MRI data. Standard vector-autoregressive models are limited by their linearity assumptions, while nonlinear general-purpose, large-scale temporal models, such as LSTM networks, typically require large amounts of training data, not always readily available in biological applications; furthermore, such models have limited interpretability. We introduce here a novel approach for learning a nonlinear differential equation model aimed at capturing brain dynamics. Specifically, we propose a variable-projection optimization approach to estimate the parameters of the multivariate (coupled) van der Pol oscillator, and demonstrate that such a model can accurately represent nonlinear dynamics of the brain data. Furthermore, in order to improve the predictive accuracy when forecasting future brain-activity time series, we use this analytical model as an unlimited source of simulated data for pretraining LSTM; such model-specific data augmentation approach consistently improves LSTM performance on both calcium and fMRI imaging data.

## 1 Introduction

Complex multi-variate nonlinear dynamical systems are abundant in nature and in society, ranging from weather to brain activity and stock market behavior. Building accurate models of such systems is highly nontrivial, and considerably more difficult than modeling linear dynamics. While nonlinear dynamical systems are extensively studied in physics, control theory and related disciplines, learning such systems from data in high-dimensional settings is difficult, and traditional machine learning approaches tend to focus on generic dynamical models, such as recurrent neural networks, rather than on domain-specific types of nonlinear dynamical models, such as, for example, van der Pol (VDP) model considered in this paper.

Our goal is to propose a model that can capture the most relevant features of a complex nonlinear dynamical system, such as brain activity in neuroimaging data. Brain activity exhibits a highly nonlinear behavior that can be oscillatory or even chaotic [23], with sharp phase transitions between different states. The simplest models that can capture these behaviors are *relaxation oscillators*. One of the most famous examples is the VDP oscillator [15], used to model a variety of problems in physics. It has also played a relevant role in neuroscience given its equivalence to the FitzHugh-Nagumo equations that were introduced as a simplified model of action potential in neurons [18, 13].

In this paper, we address two main questions. First: can we actually learn *both* hidden variables and structure parameters of the van der Pol oscillator from data, when we only observe some of the variables? There has been a lot of interest in the physics and inverse problems community in simultaneously estimating states and parameters. Most approaches for learning nonlinear dynamics and parameters avoid optimization entirely by using the unscented kalman filter [26, 31, 29] or other derivative-free dynamic inference methods [16]. Derivative-free methods have limitations — there is no convergence criteria or disciplined way to iterate them to improve estimates. Optimization-based approaches for fitting parameters and dynamics are discussed by [14], who formulate parameter identification under dynamic constraints as an ODE-constrained optimization problem. We take a similar view, and use recent insights into variable projection to develop an efficient optimization algorithm for learning the hidden states and parameters of the van der Pol (VDP). The work of [14] is focused on global strategies (e.g. multiple re-starts of well-known methods); our contribution is to develop an efficient local technique for an inexact VDP formulation.

Our main scientific question is: are such models useful to neuroscience? How well can we capture dynamics and predict temporal evolution of neural activity, and are these results interpretable in the context of prior neuroscientific knowledge? We show that the answer to those questions can be positive, but require a combination of multiple approaches, such as: (1) using both optimization and stochastic search in order to get out of potential local minima and “jump” to more promising parts of an enormous search space; and (2) using our analytical oscillatory model to pre-train generic statistical approaches, such as LSTM.

We show that the best predictive accuracy is achieved by first estimating the van der Pol model (with a relatively small number of parameters) from limited training data, and then using this model to simulate large amounts of data to pre-train a general-purpose LSTM network, pulling it to specific nonlinear dynamics, and then fine-tuning it on limited-size real data. We demonstrate that this hybrid approach consistently improves LSTM performance on both calcium and fMRI imaging data.

## 2 Calcium Imaging Data

A recently introduced technique, brain-wide calcium imaging (CaI) [11], provides for a unique perspective on neural function, recording the concentrations of calcium at sub-cellular spatial resolution across an entire vertebrate brain, and at a temporal resolution that is commensurable with the timescale of calcium dynamics [12].

In [12], light-sheet microscopy was used to record the neural activity of a whole brain of the larval zebrafish, reported by a genetically-encoded calcium marker, in vivo and at 0.8 Hz sampling rate. From the publicly available data [8] it is possible to obtain a movie of 500 frames with a 2D collapsed view of 80% of the approximately 40,000–100,000 neurons in the brain, with a resolution of 400 by 250 pixels (approximately 600 by 270 microns).

In order to obtain functionally relevant information, we performed an SVD analysis of these data ^1^; the figure 1 shows the first 5 SVD time components (left column) and the corresponding space components (right column). The spatial components show a clear neural substrate, and therefore the time components can be interpreted as traces of neuronal activity from within brain systems identified by each corresponding space components. For example, spatial components 1–5 each show pronounced but non-overlapping forebrain island-like structures, often with lateral symmetry. Moreover, the second and third spatial component include in addition the hindbrain oscillator (seen in the right panels). The corresponding second and third temporal components are dominated by oscillatory activity, consistent with the physiology of the hindbrain oscillator described in [12].

**Figure 1:**
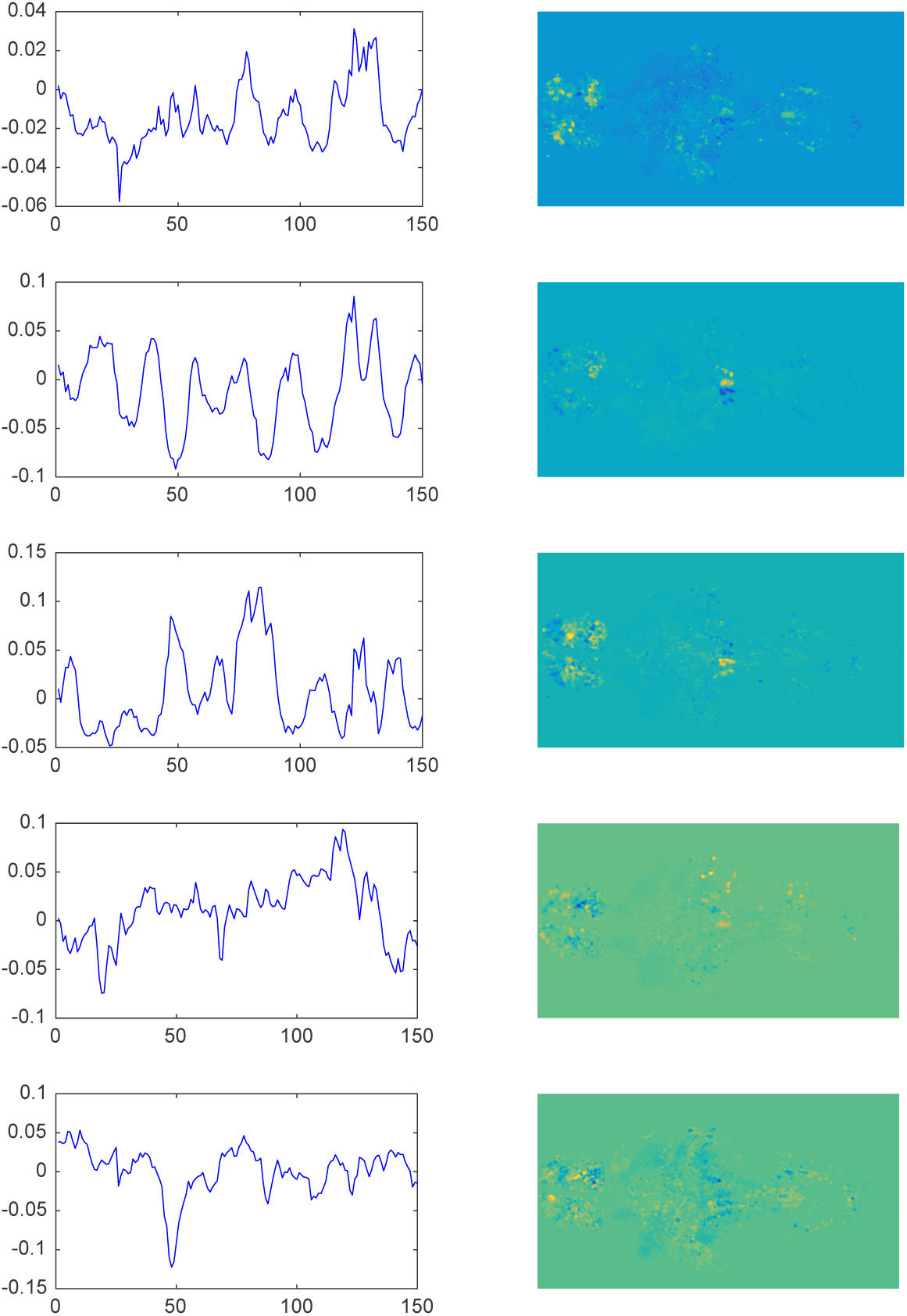
The first 5 SVD components (left column) and the corresponding space components (right column) of the zebrafish data.

## 3 Van der Pol Model of Neuronal Activity

Because neuronal calcium dynamics are largely driven by transmembrane voltage and voltage-dependent calcium channels, we model the calcium dynamics of a neuron, or small clusters of them, as a 2D differential equation with a voltage-like variable (activity), and a recovery-like variable (excitability), following similar approaches in the literature [19]. Given that one salient feature of neural systems is their propensity for oscillations, as well as sharp transitions from passive to active states, we consider the following nonlinear oscillator model for each scalar component:

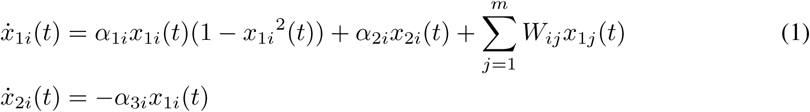

where *m* is the number of considered neural units (e.g, SVD components), *x*_1*i*_(*t*) and *x*_2*i*_(*t*) represent the (observed) activity and the (hidden) excitability variables of the *i*-th neural unit, respectively, and the *W* matrix represents the coupling strength between the observed variables, or neural units. Thus, 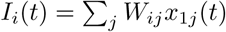 models the synaptic input to the *i*-th unit provided by other units through their observed *x*_1*j*_ variables. The parameters *α*_*ki*_ determine the bifurcation diagram of the system, allowing for a rich set of dynamical states including oscillations and spike-like responses [28, 19]. However, imaging techniques only provide information about activity *x*_1*i*_(*t*), i.e. the calcium concentration in the case of CaI. In consequence, any model-based analysis requires the inference of the excitability variable represented by hidden (unobserved) variables *x*_2*i*_.

*When the parameters α and W in* (1) *are known*, inferring the hidden components *x*_2*i*_(*t*) from observations *x*_1*i*_(*t*) is a nonlinear Kalman smoothing problem. Kalman filtering and smoothing methods are commonly used for inference on noisy dynamical systems. Since their invention [20, 21] these algorithms have become a gold standard in a range of applications, including space exploration, missile guidance systems, general tracking and navigation, and weather prediction. Optimization-based approaches with nonlinear and non-Gaussian models require iterative optimization techniques; see for example the survey of [2]. Dynamical modeling was applied to nonlinear systems early on by [1, 15]. More recently, the optimization perspective on Kalman smoothing has enabled further extensions, including inference for systems with unknown parameters [7], systems with constraints [6], and systems affected by outlier measurements for both linear [10, 24, 9] and nonlinear [3, 4] models.

Building on above perspective, we address the challenging problem of estimating from data both the parameters *α*_*ki*_, *W* and the hidden variable *x*_2*i*_(*t*). To the best of our knowledge, *this work is the first to propose an approach for learning a coupled van der Pol oscillator model from data.* We develop a method to find the hidden variables (*x*_2*i*_(*t*)) from the observed ones (*x*_1*i*_(*t*)) for given parameter settings, and to learn unknown parameter settings themselves. Indeed the problems are coupled; however, rather than using alternating optimization (closely related to EM), we use fast optimization techniques available for nonlinear Kalman smoothing to fully minimize over the hidden states for each update of the unknown parameters. The algorithm can be understood in the framework of recent results on variable projection (partial minimization), which is efficient for dealing with nonconvex, possibly ill-conditioned problems. While detailed convergence and sensitivity analysis of this algorithm is a topic of ongoing work, we present here promising results showing that the obtained van der Pol model can accurately capture nonlinear dynamics in the training data, and, furthermore, can be used to predict the future time series (test data); in addition, we show predictive performance is boosted by combining the van der Pol with LSTM networks.

## 4 Estimating van der Pol Parameters: ODE-Constrained Inference

We discretize the ODE model in equation (1), and formulate a joint inference problem for the state space *x* and parameters *α, W* that is informed by noisy direct observations of some components; and constrained by the discretized dynamics.

### Inference for a single component

For time index *k*, let 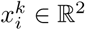 denote the *i*th component of the van der Pol model given earlier in the equation (1), so 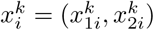, i.e. the state contains both observed and hidden variables. The discretized dynamics governing the evolution can be written

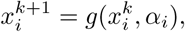

where *g* is a first-order Euler discretization of the non ear ODE (1). The *α*_*i*_ inform the evolution of the entire time series 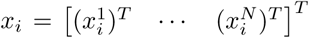. Given an initial and possibly inaccurate state 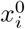, we form a vector 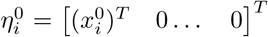, and describe the dynamics of the entire *i*th component in compact form as 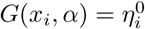, with

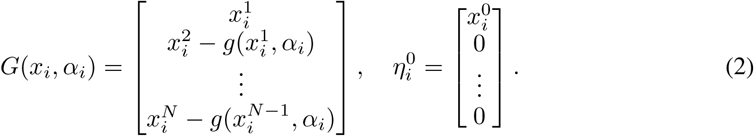

Given noisy observations

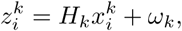

we obtain consider ODE-constrained optimization problem for the *i*th component:

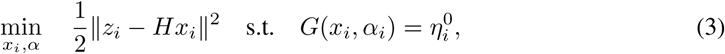

Problem (3) is challenging because (1) the ODE constraint function *G* is nonlinear in *x*_*i*_, and (2) because it is a joint optimization problem over *α*_*i*_ and *x*_*i*_. To solve this problem, we use the technique of partial minimization [5]^2^, often used in PDE-constrained optimization [30].

Rewriting (3) with a quadratic penalty, we obtain the relaxed problem

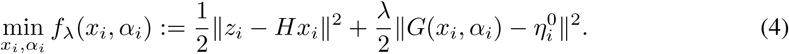

The key idea is to then use partial minimization with respect to *x*_*i*_ at each iteration of *α*_*i*_ and optimize the value function:

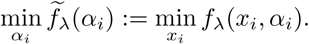

The intuitive advantages of this method (find the best state estimate for each *α* regime) are borne out by theory. In particular, for a large class of models, the objective function *f*_*λ*_(*α*_*i*_) is well-behaved for large *λ*, unlike the joint objective *f*_*λ*_(*x*_*i*_, *α*_*i*_) [5]^3^.

Evaluating *f*_*λ*_(*α*) requires a minimization routine. We compute gradient and Hessian approximations

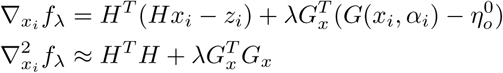

where *G*_*x*_ = *∇*_*x*_*G*(*x*_*i*_, *α*_*i*_). Evaluating *f*_*λ*_ requires obtaining an (approximate) minimizer 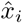. With 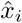 in hand, *∇*_*α*_*i f*_*λ*_ can be computed using the formula

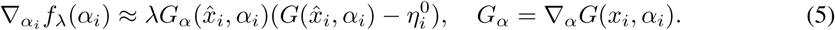

The accuracy of the inner solve in *x*_*i*_ can be increased as the optimization over *α*_*i*_ proceeds. Constraints can also be placed on *α*_*i*_ to eliminate non-physical regimes or to incorporate prior information.

**Extension to *m* components** In addition to estimating the dynamic parameters *α*, we are also interested in inferring the connectivity matrix *W*. Extending the model to *m* components, let *x* contain *m* components *x*_*i*_, so that in particular the *k*-th component *x*^*k*^ contains 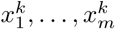; and let *α* contain *m* parameter sets *α*_*i*_. We can now write down the full nonlinear process model *G* as

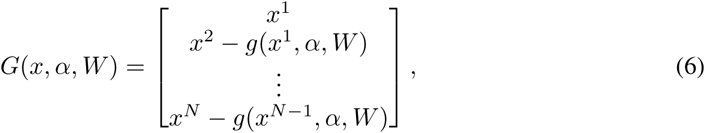

with *x∈*ℝ^2*mN*^, and the dynamics in the previous section replicated across the *m* components. Without the *W* matrix, this would be *m* independent models written jointly. The *W* adds linear coupling across the components.

The optimization approach for *m* components is analogous to the single-component case, but includes *m* components simultaneously, and also infers the coupling matrix *W*:

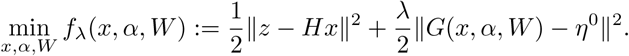

Just as for a single component, we optimize this objective using partial minimization in *x* and working with the value function

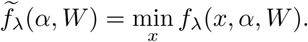

For the *m*-parameter case, we optimize over *x* at each iteration using the Gauss-Newton method detailed in the previous section. The outer iteration is a fast projected gradient method for minimizing 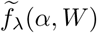 subject to simple bound constraints.

## 5 Learning van der Pol: Variable-Projection + Stochastic Search

Optimizing van der Pol model can benefit considerably from a good initialization of its parameters, as we observed in multiple experiments. To improve initialization, we start with a random walk (stochastic search) in the parameter space, aiming at producing a reasonably good starting point for the optimization procedure; given a combination of parameters, we simulate time series using the corresponding van der Pol model, and measure the correlation and the mean-squared error between the simulated and the real (training) data, discarding the parameters whose performance metrics are under some threshold. Once a sufficiently high-performing model is found, we switch to the variable-projection (VP) method described above, initialized with the current parameters, which are now optimized even further (Figure 2). The whole process of alternating between stochastic search and VP optimization is repeated several times, since, consistently with reported works [14, 27], a hybrid stochastic-deterministic method performs better than a sole local optimization method for complex problems. This combined procedure will be referred to in our subsequent section as simply *van der Pol optimization*.

**Figure 2:**
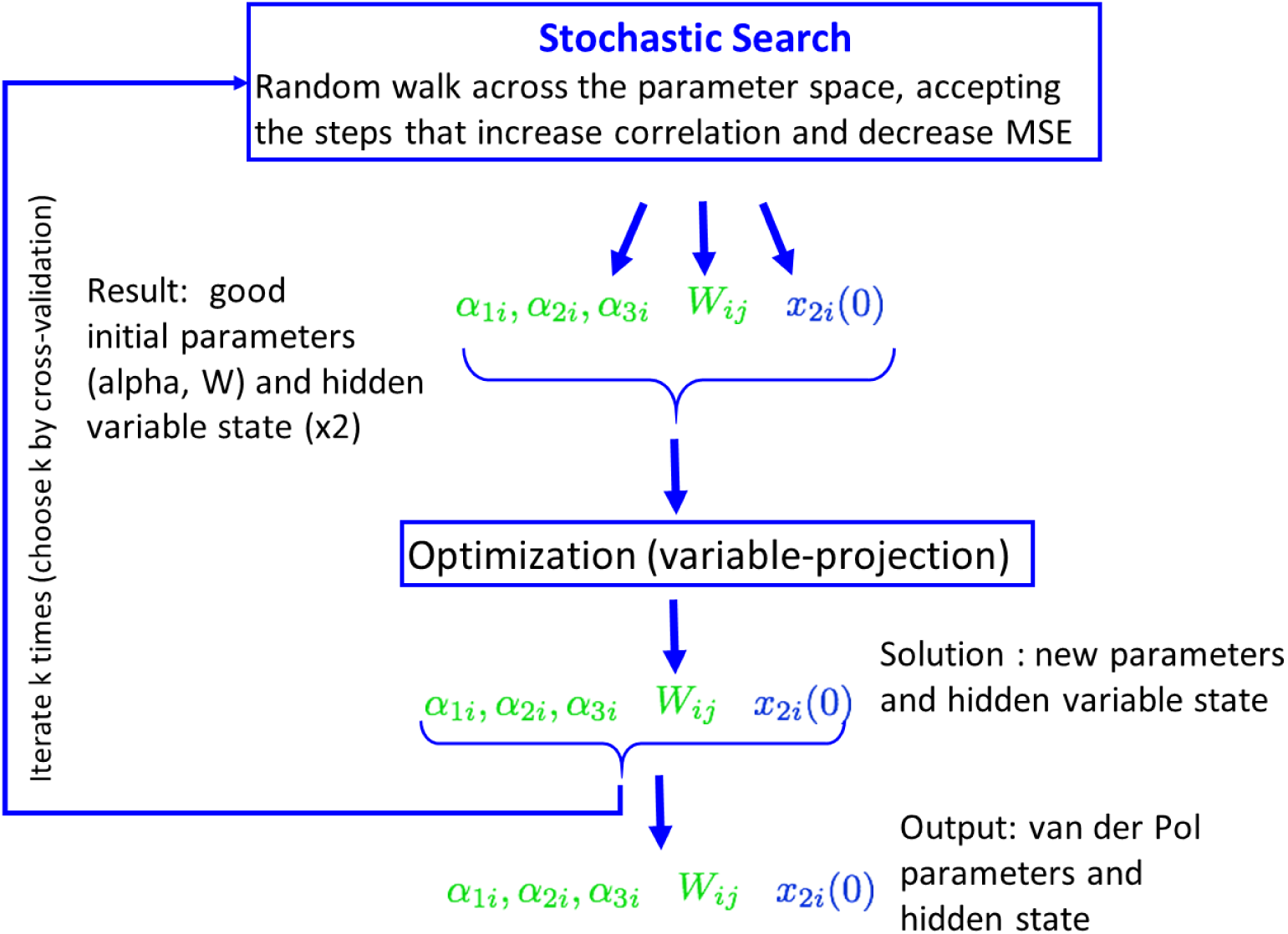
Van der Pol optimization procedure: variable-projection augmented with stochastic search.

### Stochastic search: implementation details

We start with an initial guess for *α*_*ki*_ (same value for all univariate oscillators), zero-connectivity matrix *W* and a random guess for the initial condition of the hidden variables, *x*_2*i*_(0). At every stochastic search step, these parameters are updated as described below, and the differential equations with the new parameters are integrated; if the resulting time-series solution improves the fit to the training data, the new parameters are accepted, otherwise they are dismissed. As a measure of the goodness of fit we use a linear combination of the Symmetric Mean Absolute Percentage Error and the Pearson correlation. In the first stage of our search, we only update *α*_*ki*_ and *x*_2*i*_(0), while keeping zero weight matrix *W* (i.e., disconnected components). In each step, one of the components *x*_*i*_ is chosen randomly, and its corresponding *α*s and *x*_2*i*_(0) are changed using a Gaussian random walk. Steps with larger variance are taken infrequently to escape potential local minima. After this initialization, we update all parameters including *W*. All components of *W* change at every (low-variance) random step.

### VP+Stochastic Search: implementation details

We use up to 200000 stochastic steps with a maximum of 50 outer iterations of the VP optimization for every 1000 stochastic steps. The *W* matrix is held to zero for the first ‘burn-in’ 15000 iterations. Large stochastic steps are performed after every 30 small steps; the variance in *W* steps increases by one order of magnitude (from 0.01 to 0.1); for *α* steps, the variances remains the same (0.1), but larger steps involve changing all components at once, rather than one at a time in smaller steps. We use *λ* = 3*e*9 in the VP optimization formulation.

### Time series prediction

Once a van der Pol model is trained on a given time window, we can use it to predict the future time series, by integrating the model with the given parameters and the initial hidden state variable.

### Interpretability

Note that one of the advantages of the analytical van der Pol model is its interpretability, as it learns the interaction matrix *W* among different spatial components, i.e. brain areas.

## 6 Hybrid approach: vdP-LSTM

An alternative to learning an analytical model, such as van der Pol, is to use some generic method for time-series prediction, such as, for example, recurrent neural networks, e.g., LSTM. Herein, we used the classical LSTM model proposed by [17], a popular extension of Recurrent Neural Network (RNN) models with improved memory. Our LSTM networks, implemented in Keras, contained two layers, 128 units in each layer, followed by the fully-connected layer and linear activation; we used the mean squared error and the optimizer RMSProp; the drop-out rate was set to 0.8. We used LSTM for multivariate time-series prediction, where each time point *t* is represented by an *n*-dimensional vector (corresponding, in our case, to temporal components of the data at time *t*). We denote by *LSTM* (*k*) the model which uses the previous *k* time points to predict the *k* + 1-st time point. The prediction of the time step *k* + 2 is performed by shifting the window of length *k* one step forward and using the prediction for the *k* + 1-st data point a new data point, iteratively. In our experiments, several values of *k* were tried and *k* = 6 was selected.

### vdP-LSTM: LSTM Pretrained with vdP Simulations

Training LSTMs requires a large number of samples, while our data were limited to only 500 time points (including training and test subsets). On the other hand, given our prior knowledge about the data, such as nonlinear dynamic behavior of certain type (e.g., van der Pol with specific parameters), one can hypothesize that providing LSTM with information about such domain-specific dynamics can potentially improve its performance.

Thus, we propose a simple approach for providing general-purpose LSTM with prior information about the domain-specific dynamics, namely, a data-augmentation approach which pretrains LSTM on a large amount of simulated data obtained from a fitted van der Pol model, before fine-tuning LSTM on a relatively small amount of available real data. Such pretraining on the data simulated from our analytical model serves as a regularizer in the absence of large training data sets, biasing LSTM towards the type of dynamics we expect in the data. Training the van der Pol on the same amount of data is easier, since there are far fewer parameters to be estimated than for a typical LSTM.

### vdP-LSTM: implementation details

We train *n* = 18 van der Pol models on the training data; for each of those models, we simulate *m* = 15 noisy versions of time series (each of length *k* = 160) obtained by integrating each model; we take *m* = 15, *n* = 18, and *k* = 160.

LSTM was pretrained with 100 epochs using the above simulated data, and then trained with 50 epochs on the real training dataset; the number of epochs was selected so that the total number of samples used for training was the same for both simulated and real training data.

## 7 Experiments

We now present our empirical results, including (1) van der Pol model fit on training data; (2) predictive accuracy when forecasting time series using van der Pol, LSTM and hybrid vdP-LSTM, with a linear Vector Auto-Regressive (VAR) model used as a baseline; (3) a brief discussion of interpreting the van der Pol interaction matrix.

### Evaluating van der Pol model fit on real data

We evaluated multiple runs of van der Pol estimation procedure described above, combining stochastic search with VP optimization. Figure 4 shows the fit to the training data achieved by one of the best-performing model; the correlations between the actual data and the model predictions are high, ranging from 0.76 to 0.83 for all six components.

**Figure 3:**
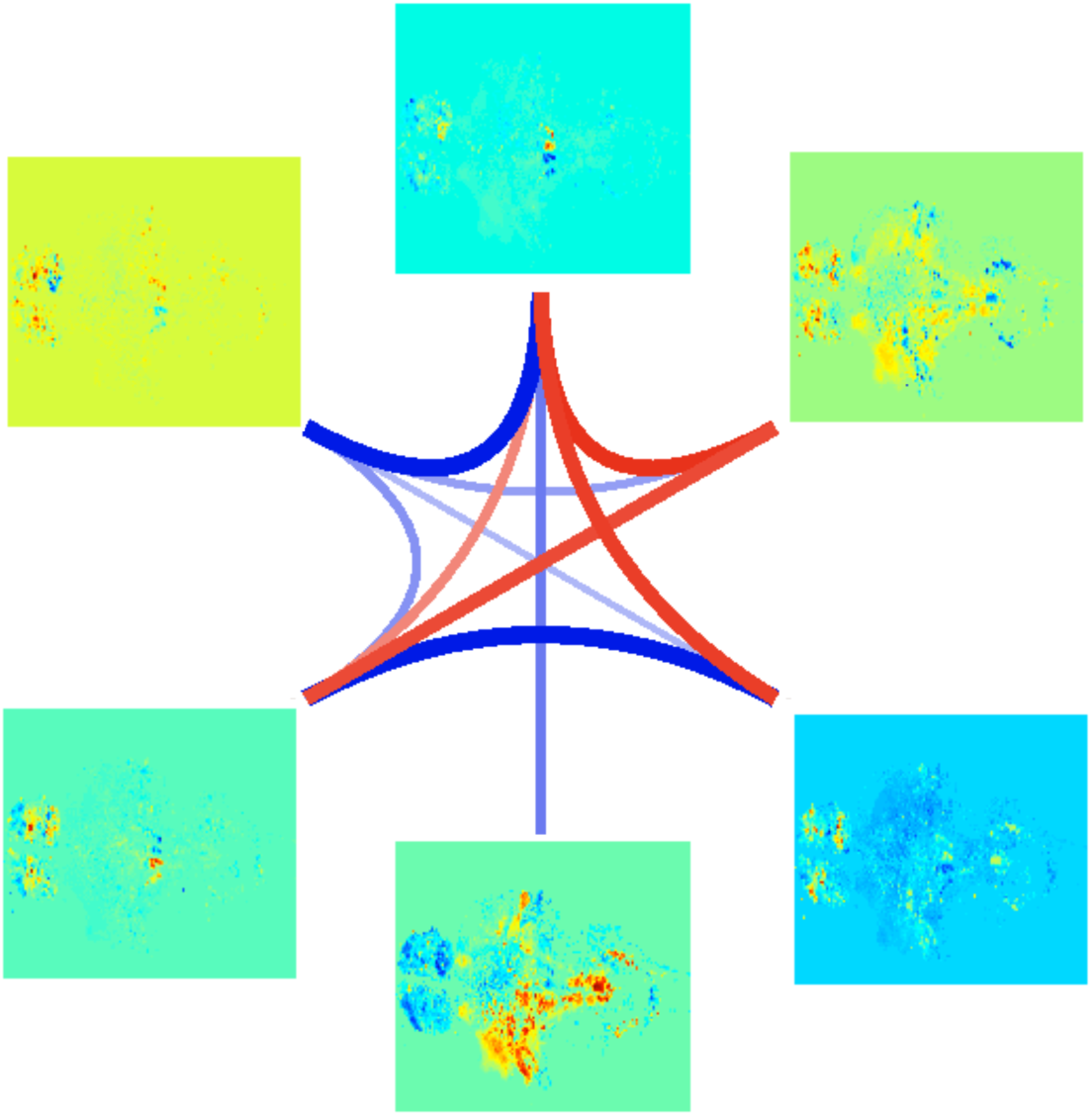
Spyder graph representing the strongest links between the spatial support of the components, as interpreted from *W*. Red represent negative and blue, positive.

**Figure 4:**
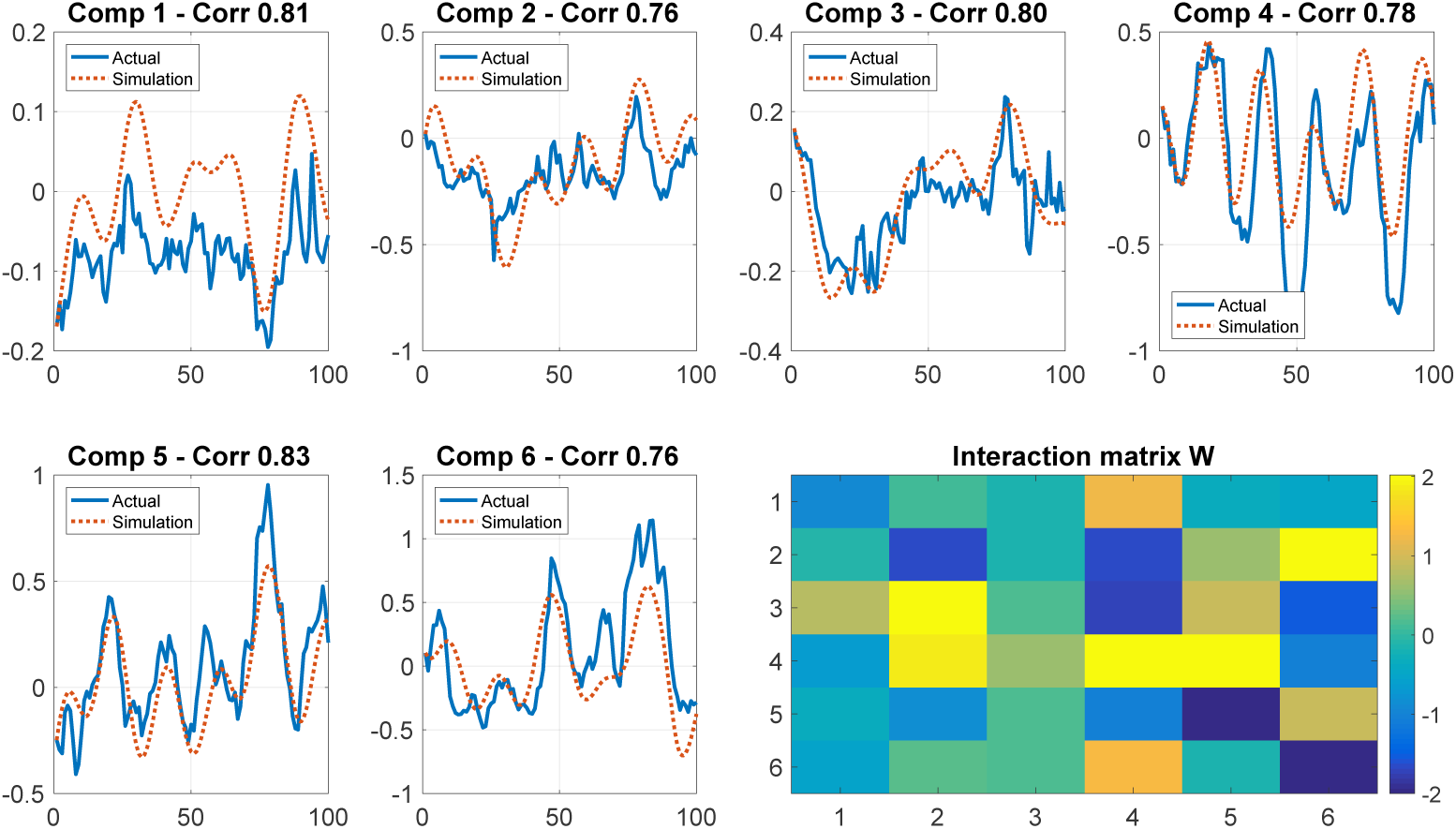
Van der Pol model fit on training data; correlations between the true and predicted time series for each of the six temporal SVD components. Bottom-right: an interaction matrix *W*.

### Interpretability

In the bottom right corner of the Figure 4, we plot the coupling matrix *W*. We interpret the entries of W as the effective connectivity between the spatial support of the components. Thus, *W* contains interesting information about interactions (positive and negative) across different brain regions/subnetworks. For example, we observe a strong positive interactions between the components 4 and 5, which correspond to the brain areas where an “flip-flop” oscillating behavior can be clearly observed (e.g., see the 2D version of the temporal data at https://youtu.be/lppAwkek6DI). Figure 3 presents a spyder graph representing the strongest links between the spatial support of the components, obtained from *W*. Using current knowledge of zebrafish and human neuroanatomy, it is possible to validate to what extent this effective connectivity (at least in absolute value) is consistent with real neural tracts.

### Prediction on test data

Figures 5 and 6 show the median correlation between the true and predicted values, and the root-means square error, respectively, for several predictive methods: vector autoregressive (VAR) model (red), van der Pol model (green), LSTM (blue), and vdP-LSTM, i.e. LSTM pretrained on the data simulated using the above van der Pol model (orange). Here we estimated parameters of the models on 100 consecutive points of training data, and then predicted the next 30 points (x-axis plots the index of the time points being predicted). Shaded area around each curve represents the standard error. The linear VAR model (red) performs poorly, unable to capture the nonlinear dynamics; van der Pol (green) outperforms LSTM (blue) in the beginning, but then LSTM catches up; the hybrid vdP-LSTM model combines the best of both. Similarly, the hybrid approach performs best in terms of RMSE error (Figure 6).

**Figure 5:**
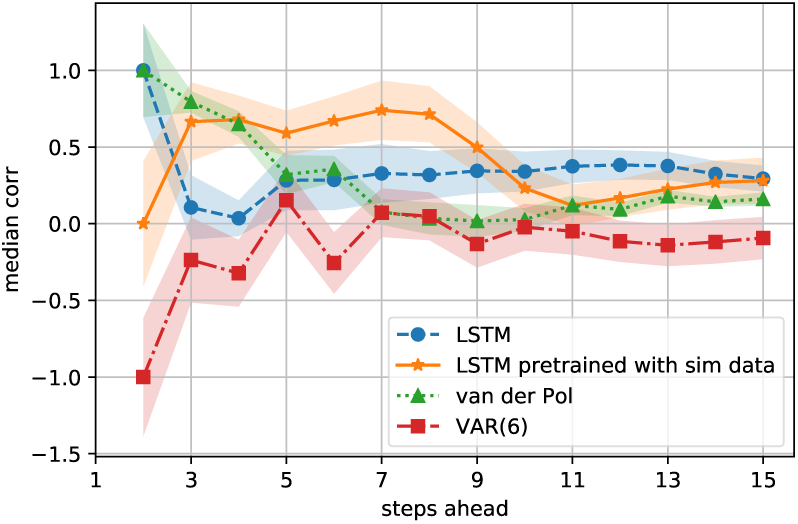
Calcium imaging: predictive performance measured by correlation between the actual and predicted time series.

**Figure 6:**
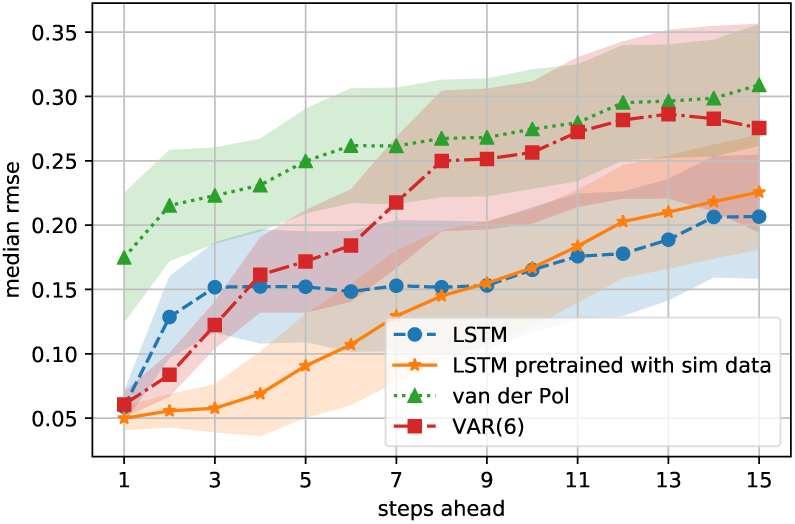
Calcium imaging: predictive performance measured by the root mean square error (RMSE) between the actual and predicted time series.

### 7.1 Functional MRI Data

We also tested our approach on a functional MRI (fMRI) and obtained promising preliminary results. Though VP optimization was not yet applied on top of the stochastic search (experiments are in progress), we already obtained results similar to the ones seen on calcium data. We used resting-state fMRI data from 10 healthy control subjects, obtained from the Track-On HD dataset [22]. For each subject we had two datasets corresponding to two different visits. We used 15 ICA components, 160 time points each. The datasets from the first visit were used for training, and the ones from the second visit were used for testing. For each training dataset, we ran stochastic search 10 times, and from each run used 50 models which correlated highest with the training data for subsequent simulations and LSTM pre-training; i.e., for each subject, we simulated 15 coupled time series, each of length 160, from 500 different (but related) van der Pol models.

In addition, for comparison with a standard method of data augmentation, 500 noisy datasets (i.e., multivariate time series, with 15 components), also of 160 time steps, were created from each subject’s training dataset by adding Gaussian noise with mean 0 and standard deviation 0.1 to the normalized real data.

The LSTM architecture used was the same as for the calcium imaging experiments. Each subject was trained separately, as a different instance of the experiment. Each dataset contained 15 time series, with 160 time points each. LSTM was trained with 15 epochs. The data-augmented LSTMs were first trained either with the noisy datasets or with the van der Pol-simulated data described above for 15 epochs followed by the training with 15 epochs of real training data (the first visit data for a given subject).

Figures 7 and 8 summarize the correlation and RMSE performance, respectively, of several methods we tried on fMRI data, such as VAR, LSTM, as well as LSTM pretrained with noisy version of real data (standard data-augmentation approach), and vdP-LSTM (LSTM pretrained on van der Pol simulated data). We see much smaller standard error (shaded area around the plots), due to larger number of experiments per point (more fMRI data). Overall, we clearly see that VAR performs poorly, and vdP-LSTM, augmented with simulated data, outperforms both LSTM, and LSTM augmented with noisy data, in terms of correlation and RMSE.

**Figure 7:**
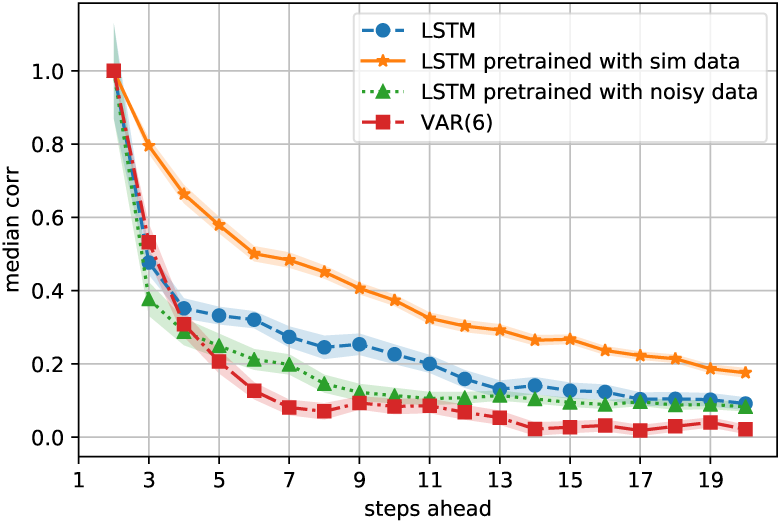
Functional MRI: predictive performance measured by correlation between the actual and predicted time series.

**Figure 8:**
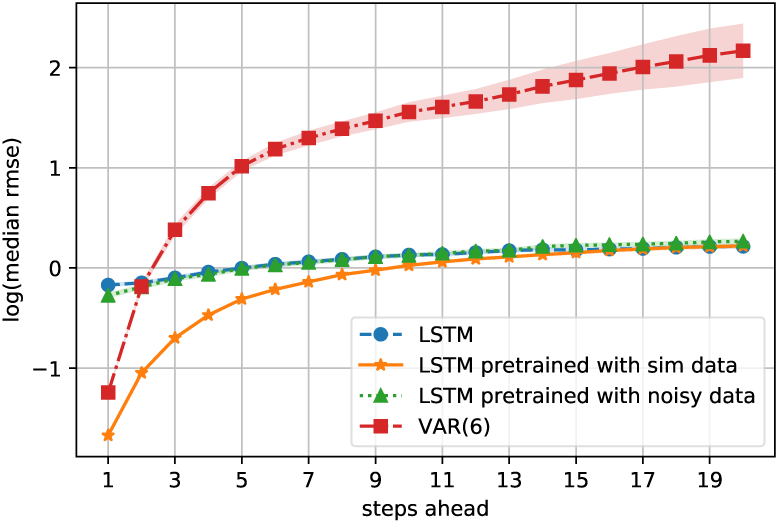
Functional MRI: predictive performance measured by the root mean square error (RMSE) between the actual and predicted time series.

## 8 Conclusions

Motivated by the challenging problem of modeling nonlinear dynamics of brain activations in calcium imaging, we propose a new approach for learning a nonlinear differential equation model: a variable-projection optimization approach to estimate the parameters of the multivariate coupled van der Pol oscillator. We show how to learn this nonlinear dynamical model, and demonstrate that it can accurately capture nonlinear dynamics of the brain data. Furthermore, in order to improve the predictive accuracy when forecasting future brain activity, we used the learned van der Pol to pretrain LSTM networks, thus imposing an oscillator prior on LSTM; the resulting approach achieves highest predictive accuracy among all methods we evaluated.

## Footnotes

1 There are multiple alternative approaches to feature extraction/representation learning and dimensionality reduction, which can be explored in this setting, including other component analysis methods (NMF, ICA), sparse coding/dictionary learning, and various autoencoders. However, before diving into more complex feature extraction, we would like to develop an approach to modeling a coupled dynamical system, which is a nontrivial task even with a relatively small number of SVD components.

2 For particular instances, partial minimization is often called *variable projection*.

3 The Lipschitz constant of the gradient of 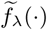 stays bounded as *λ → ∞*, which is clearly false for *f*_*λ*_(*·,·*).

